# A mitochondrial long-chain fatty acid oxidation defect leads to uncharged tRNA accumulation and activation of the integrated stress response in the mouse heart

**DOI:** 10.1101/2021.05.13.443905

**Authors:** Pablo Ranea-Robles, Natalya N. Pavlova, Aaron Bender, Andrea S. Pereyra, Jessica M. Ellis, Brandon Stauffer, Chunli Yu, Craig B. Thompson, Carmen Argmann, Michelle Puchowicz, Sander M. Houten

## Abstract

The heart relies mainly on mitochondrial fatty acid β-oxidation (FAO) for its high energy requirements. Cardiomyopathy and arrhythmias can be severe complications in patients with inherited defects in mitochondrial long-chain FAO, reinforcing the importance of FAO for cardiac health. However, the pathophysiological mechanisms that underlie the cardiac abnormalities in long-chain FAO disorders remain largely unknown. Here, we investigated the cardiac transcriptional adaptations to the FAO defect in the long-chain acyl-CoA dehydrogenase (LCAD) knockout (KO) mouse. We found a prominent activation of the integrated stress response (ISR) mediated by the eIF2α/ATF4 axis in both fed and fasted states, accompanied by a reduction in cardiac protein synthesis during a short period of food withdrawal. Notably, we found an accumulation of uncharged tRNAs in LCAD KO hearts, consistent with a reduced availability of cardiac amino acids, in particular, glutamine. We replicated the activation of the cardiac ISR in hearts of mice with a muscle-specific deletion of carnitine palmitoyltransferase 2 deletion (*Cpt2*^M-/-^). Our results show that perturbations in amino acid metabolism caused by long-chain FAO deficiency impact cardiac metabolic signaling, in particular the ISR, and may play a role in the associated cardiac pathology.

## Introduction

The adult healthy heart is particularly reliant on mitochondrial fatty acid oxidation (FAO) for its energetic needs when compared to other organs (Allard et al. 1994). A recent comprehensive study reported that fatty acids provide around 85% of the myocardial ATP needed in healthy human subjects (Murashige et al. 2020). Pathological hypertrophic cardiac growth is associated with a reprogramming of heart metabolism, characterized by decreased fatty acid metabolism and increased reliance on glucose metabolism. This cardiac metabolic remodeling has been described in animal models (Allard et al. 1994; Aubert et al. 2016; Lai et al. 2014; Osorio et al. 2002; Ritterhoff et al. 2019) and human patients (Bedi et al. 2016; Dávila-Román et al. 2002; Gibb & Hill 2018; Murashige et al. 2020; Sack et al. 1996). The importance of FAO in cardiac health is supported by the observation that patients with an inherited disorder of FAO may develop hypertrophic cardiomyopathy and arrhythmias (Bonnet et al. 1999; Kelly & Strauss 1994; Knottnerus et al. 2020). Although altered myocardial fatty acid metabolism has been documented in patients with FAO defects (Bergmann et al. 2001; Kelly et al. 1993), the metabolic and transcriptional adaptations that underlie the cardiac pathophysiology remain largely unknown.

Mouse models are useful to investigate the effects of FAO deficiency on cardiac metabolism and function. The long-chain acyl-CoA dehydrogenase (LCAD) KO (*Acadl*^-/-^) mouse serves as a good model for human long-chain FAO disorders such as very long-chain acyl-CoA dehydrogenase deficiency (VLCAD) and carnitine palmitoyltransferase 2 (CPT2) deficiency. The LCAD KO mouse recapitulates prominent symptoms of VLCAD deficiency including fasting-induced hypoketotic hypoglycemia and cardiac hypertrophy (Bakermans et al. 2011; Chegary et al. 2009; Cox et al. 2009; Houten et al. 2013; Kurtz et al. 1998). Previous studies have revealed a fasting-induced cardiac dysfunction in LCAD KO mice, characterized by an impaired left ventricular performance including an increase in end systolic volume, and reduced ejection fraction, stroke volume, cardiac output, peak ejection rate and peak late filling rate (Bakermans et al. 2011, 2013). At the metabolic level, we previously described an impairment of amino acid metabolism in LCAD KO mice causing a shortage in the supply of gluconeogenic precursors (Houten et al. 2013) and an unmet anaplerotic need in the heart of fasted LCAD KO mice (Bakermans et al. 2013). These alterations are characterized by increased cardiac glucose uptake and oxidation, suppressed glucose-alanine cycle, decreased circulating levels of gluconeogenic amino acids, decreased level of cardiac TCA cycle intermediates and impaired protein mobilization (Bakermans et al. 2011, 2013; Houten et al. 2013). The molecular responses of the FAO-deficient heart to these metabolic challenges are unknown, but could shed some light into the mechanisms underlying the associated cardiac pathology.

The integrated stress response (ISR) is a well-conserved stress response that is orchestrated by phosphorylation of the alpha subunit of the eukaryotic translation initiation factor 2 complex (eIF2α) (Harding et al. 2000). Different stressors are sensed by four specialized eIF2α kinases (HRI, PKR, PERK, GCN2) leading to phosphorylation of S51 on eIF2α (Costa-Mattioli & Walter 2020). Phosphorylated eIF2α blocks the eIF2’s guanine nucleotide exchange factor eIF2B and results in a reduction of protein synthesis. At the same time, the ISR promotes the selective translation of a pool of mRNAs that contain short inhibitory upstream open reading frames in their 5’ UTR, including activating transcription factor 4 (ATF4) (Vattem & Wek 2004). ATF4 target genes encode, among others, amino acid biosynthesis enzymes and amino acid transporters, as part of the cellular response specific to amino acid depletion (Harding et al. 2003; Noland et al. 2007).

Here, we used transcriptome analysis to identify molecular changes in the heart in response to a long-chain FAO deficiency. To identify early and late events, we also characterized the development of changes in metabolite levels and metabolic signaling during an overnight food withdrawal. We identify an activation of the ISR in the FAO-deficient heart that is largely independent of the feeding status.

## Methods

### Animal experiments

LCAD KO (*Acadl*^-/-^) mice (Kurtz et al. 1998) on a pure C57BL/6 background were maintained on a normal chow diet (PicoLab Rodent Diet 20, LabDiet). The *Acadl* allele was genotyped using an SNP-based assay (Luther et al. 2012). Cardiac and skeletal muscle CPT2-deficient mice (*Cpt2*^M-/-^) were generated as previously described (Pereyra et al. 2017). Briefly, a loxP flanked specific sequence of the *Cpt2*^M-/-^ gene was excised by the Cre recombinase under the control of the muscle creatine kinase promoter. Littermates lacking Cre enzyme expression were used as controls. *Cpt2*^M-/-^ mice and controls were maintained on a custom-made control diet (Envigo TD 94045) (Pereyra et al. 2021). Mice were housed in pathogen-free rooms with a 12 h light/dark cycle. All animal experiments were approved by the IACUC of the Icahn School of Medicine at Mount Sinai (# IACUC-2014-0100) and the Purdue University IACUC (assurance #A3231-01), and comply with the National Institutes of Health guide for the care and use of Laboratory animals (NIH Publications No. 8023, 8th edition, 2011).

There were six experimental cohorts of adult mice used in this study. The cohort for the RNA-sequencing experiments consisted of four WT and four LCAD KO male mice (Diekman et al. 2014). There was a second cohort to validate the RNA-sequencing results by immunoblot with six WT and seven LCAD KO 10-month-old male mice (retired breeders). The cohort for the time course analysis after food withdrawal consisted of 36 WT (18 males and 18 females) and 35 LCAD KO (17 males and 18 females) 3-8 month-old mice. The cohort used for protein synthesis measurement consisted of 12 WT (9 males and 3 females) and 15 LCAD KO (11 males and 4 females). The cohort for tRNA charging and amino acid measurement after 4 hours of food withdrawal consisted of 12 WT (5 males and 7 females) and 8 LCAD KO (5 males and 3 females) mice. The sixth cohort consisted of 6 control (3 males and 3 females) and 6 *Cpt2*^M-/-^ KO (3 males and 3 females) 7-week-old mice.

For fractional protein synthesis (FSR, %), as measured by the ^2^H_2_O method (Bederman et al. 2006, 2015; Gasier et al. 2010; Yuan et al. 2008), animals were injected with deuterium oxide (^2^H_2_O) as 0.9% NaCl either at 7 a.m. (fed and morning food withdrawal conditions) or at 6 p.m. (the beginning of the active phase for mice). Food withdrawal was performed by removing the food and placing the mice in a clean cage. We aimed to reach 4% enrichment of total body water using the following formula: *Volume of heavy water injected (mL) = Body weight (g) * 0.75 (Water content in the body) * 0.04 (% of enrichment)*, as previously described (Ranea-Robles et al. 2021). After 7 hours, a small amount of blood was drawn from the animals to measure ^2^H_2_O enrichment, and immediately after, the mice were euthanized by exposure to CO_2_. The heart was quickly frozen with a freeze-clamp, wrapped in aluminum foil, temporarily stored in liquid nitrogen, and ultimately transferred and stored at −80°C.

Blood glucose was measured after overnight food withdrawal using Bayer Contour blood glucose strips. Mice were euthanized by exposure to CO_2_ except mice from the time course cohort, which were euthanized using pentobarbital (intraperitoneal 150 mg/kg in 0.9% NaCl). Blood was collected from the inferior vena cava for the preparation of EDTA plasma. Organs were snap frozen in liquid nitrogen and stored at −80°C.

### RNA-seq, differential gene expression, pathway enrichment

RNA was isolated using QIAzol lysis reagent followed by purification using the RNeasy kit (Qiagen). RNA samples were submitted to the Genomics Core Facility at the Icahn Institute and Department of Genetics and Genomic Sciences, and sequenced essentially as previously described (Argmann et al. 2017). Differential gene expression analysis was conducted with R packages DESeq2 (Love et al. 2014) as previously described (Argmann et al. 2017). Cut-off for differential gene expression was chosen at an adjusted p-value (Benjamini-Hochberg) of < 0.05.

Pathway enrichment analysis was performed using the ClueGo v2.5.6 and CluePedia v1.5.6 plugins in Cytoscape (v. 3.7.2) with the GO (BP), KEGG, Wiki and Reactome databases (02/2020 download) (Bindea et al. 2009, 2013). Lists of genes, either up-or down-regulated (@logFC > 0.58, adjP < 0.05) in LCAD KO vs control hearts were queried as two separated gene clusters. Pathway enrichment was also performed on the up-regulated genes shared between the LCAD KO heart (@logFC > 0.58, adjP < 0.05) and the muscle-specific CPT2 KO heart (*Cpt2*^M-/-^ mice, all 3 methods, adjP < 0.05) transcriptional signatures. The *Cpt2*^M-/-^ heart transcriptional signature was curated from Pereyra et al. (Pereyra et al. 2017). Pathway enrichment analysis with this dataset was performed with the KEGG, Wiki and Reactome databases. Cut-off for pathway enrichment was chosen at an adjusted p value (Bonferroni step down) of < 0.05 and the minimum number of Genes/pathway selection = 5. *Cpt2*^M-/-^ heart up-and down-regulated genes were tested for enrichment in up- or down-regulated DEGs (at adj p < 0.05) between LCAD and WT hearts using the Fisher’s exact test (FET) and p values were adjusted using BH procedure.

iRegulon v1.3 was used to predict transcriptional regulators of heart differentially expressed genes (Janky et al. 2014). Input genes included genes up-regulated (@logFC > 0.58, adjP < 0.05) in LCAD KO vs control hearts, and the shared up-regulated genes between the LCAD KO heart and the *Cpt2*^M-/-^ heart transcriptional signatures. Input genes were converted from mouse to human orthologs using g:Orth from g:Profiler (Raudvere et al. 2019). The following options *Motif collection*: 10K (9713 PWMs), *Putative regulatory region*: 10kb centered around TSS (10 species) alongside the program default settings for *Recovery* and *TF prediction* options were selected for the analysis.

### Immunoblot analysis

Lyophilized mouse hearts were homogenized in RIPA buffer supplemented with protease and phosphatase inhibitors (Pierce) using a TissueLyser II apparatus (Qiagen), followed by sonication and centrifugation (10 min at 12,000 rpm at 4°C). Total protein concentration was determined using the BCA method. Proteins were separated on Bolt 4-12% Bis-Tris Plus gels (Invitrogen, Thermo Fisher Scientific Inc.) and blotted onto nitrocellulose (926-31092, LI-COR) or PVDF (IPFL00010, Millipore) membranes using a semi-dry transfer system (TE77X, Hoefer). Equal gel loading was verified by Ponceau S staining. Proteins were detected using the following primary antibodies: anti-phospho-eIF2α (Ser51, 9721, Cell Signaling), anti-eIF2α (9722, Cell Signaling), anti-ATF4 (11815, Cell Signaling), anti-c-Myc (5605, Cell Signaling), anti-ASNS (sc-365809, Santa Cruz), anti-phospho-4E-BP1 (Thr37/46, 2855, Cell Signaling), anti-4E-BP1 (9644, Cell Signaling), anti-SQSTM1/p62 (5114, Cell Signaling), and anti-LC3B (3868, Cell Signaling). PVDF membranes were used to detect LC3B bands. The following secondary antibodies were used: goat anti-mouse and goat anti-rabbit secondary antibodies IRDye 800CW and IRDye 680RD (926-32 210, 926-68 070, 926-32 211, 926-68 071) from LI-COR, and Alexa Fluor AffiniPure Goat Anti-Mouse IgG light chain specific secondary antibodies 790nm and 680 nm (115-655-174 and 115-625-174) from Jackson ImmunoResearch Inc. (West Grove, Pennsylvania). Proteins were visualized in an Odyssey CLx Imager (LI-COR). Band intensity was quantified using the Fiji distribution of ImageJ 1.x (Schindelin et al. 2012; Schneider et al. 2012). For some blots, HEK-293 cells were cultured and treated with 2 µg/mL of tunicamycin (ab120296, Abcam) for 6 hours. Protein was extracted and used as positive control for the identification of p-PERK (T980), PERK, and ATF6.

### Metabolite analysis

Plasma acylcarnitines and amino acids were measured and analyzed by liquid chromatography and tandem mass spectrometry (LC-MS/MS) on an Agilent 6460 Triple Quadrupole Mass Spectrometer by the Mount Sinai Biochemical Genetic Testing Lab (now Sema4), as previously described (Le et al. 2014; Leandro et al. 2019; Ranea-Robles et al. 2020). The L-carnitine internal standard (^2^H_9_-carnitine) and the acylcarnitine internal standard mix containing ^2^H_9_-carnitine, ^2^H_3_-acetylcarnitine (C2), ^2^H_3_-propionylcarnitine (C3), ^2^H_3_-butyrylcarnitine (C4), ^2^H_9_-isovalerylcarnitine (C5), ^2^H_3_-octanoylcarnitine (C8), ^2^H_9_-myristoylcarnitine (C14), and ^2^H_3_-palmitoylcarnitine (C16), were from Cambridge Isotope Laboratories (Tewksbury, MA, USA).

### tRNA charging

Charging of tRNA was measured essentially as described (Pavlova et al. 2020). Mouse hearts for this experiment were freeze-clamped directly after euthanasia and stored at −80C. Hearts were pulverized on liquid nitrogen using the BioPulverizer (BioSpec, 59013N) and then lysed in QIAzol lysis reagent (Qiagen) using a TissueLyser II apparatus (Qiagen), followed by centrifugation (10 min at 12,000 x g at 4°C). Lysates were shaken with chloroform at a 5:1 volume ratio, centrifuged at 18,600g and the upper phase was precipitated overnight with 2.7x g of GlycoBlue (AM9515, ThermoFisher). Samples wereμ resuspended in 0.3M acetate buffer (pH=4.5) + 10 mM EDTA and reprecipitated with EtOH again. Next day, samples were resuspended in 10 mM acetate buffer + 1 mM EDTA. Of each RNA sample, 2 μg were treated with 10 mM of either sodium periodate (“oxidized”) or with sodium chloride (“non-oxidized”) and incubated for 20 min at room temperature in the dark, following by quenching with glucose for 15 min. Yeast tRNA^Phe^ (R4018, Sigma-Aldrich) spike-in control was added to each sample, after which samples were precipitated with EtOH. Samples were then resuspended in 50 mM Tris buffer (pH=9) and incubated for 50 min at 37°C, quenched with acetate buffer and precipitated. Finally, samples were resuspended in RNAse-free water and ligated to a 5’-adenylated DNA adaptor (5′-/5rApp/TGGAATTCTCGGGTGCCAAGG/3ddC/-3′ (M0373, New England Biolabs), for 3 hours at room temperature, according to (Loayza-Puch et al. 2016). Reverse transcription was performed with SuperScript IV reverse transcriptase (18090050, Thermo Scientific) according to the manufacturer’s instructions, with a primer complementary to the DNA adaptor. qPCR was performed with tRNA isodecoder-specific primers (**Table S5**) designed such that the forward (FW) primer was complementary to the 5’ end of the tRNA, and the reverse (RV) primer spanned the junction between the 3’ end of the tRNA and the ligated adaptor. Primers were designed using reference tRNA sequences from GtRNA database (http://gtrnadb.ucsc.edu/) (Chan & Lowe 2016). Ct values obtained with primers specific for yeast tRNA^Phe^ primers were subtracted from Ct values obtained with primers specific for each isodecoder of interest. The “charged fraction” value was calculated from a difference between dCt values from a non-oxidized (representing total) and oxidized (representing charged) samples for each pair.

### 2H labeling of protein-bound alanine to determine fractional protein synthesis (FSR)

^2^H enrichment of protein-bound alanine was determined on its methyl-8 derivative, as previously described (Bederman et al. 2006, 2015; Gasier et al. 2010; Yuan et al. 2008). The rate(s) of protein synthesis were calculated using the formula:

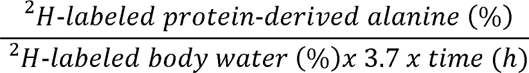

where the factor 3.7 represents an incomplete exchange of ^2^H between body water and alanine; i.e., 3.7 of the 4 carbon-bound hydrogens of alanine exchange with water. This formula assumes that the ^2^H labeling in body water equilibrates with free alanine more rapidly than alanine is incorporated into newly made protein and that protein synthesis is linear over the study.

### Statistics

We used unpaired Student t-test to analyze differences between WT and LCAD KO animals in Fig. 2 and between vehicle and L-AC in Fig. 4. We analyzed the effect of genotype (G), time (T), and their interaction on blood glucose, plasma metabolites and protein levels in the time course experiment using two-way ANOVA. *P ≤ 0.05, **P ≤ 0.01, ***P ≤ 0.001. GraphPad Prism 9 was used to compute statistical values.

## Results

### Transcriptional activation of amino acid metabolic pathways and ATF4 target genes in the heart of LCAD KO mice

To gain more insight into transcriptional adaptations of the heart to long-chain FAO deficiency, we performed RNA-sequencing analysis of WT and LCAD KO mouse heart samples collected after overnight food withdrawal and determined differentially expressed genes (DEGs). With an adjusted p value cutoff of 0.05 and a fold-change cutoff of 1.5, we obtained 2,334 DEGs (1,032 up and 1,302 down) demonstrating a large impact of FAO deficiency on the mouse heart (**Table S1A**). Pathway enrichment analysis of the DEGs revealed up-regulation of genes involved in “rRNA metabolic process”, “carboxylic acid metabolic process”, and “Hypertrophy Model” (adj p<0.05), among other pathways (Fig. 1A **and Table S1B**). Consistent with previously reported changes in amino acid metabolism in the LCAD KO mouse (Bakermans et al. 2013; Houten et al. 2013), multiple pathways related to amino acid metabolism were also enriched in the DEGs, including “cellular amino acid metabolic process” and “amino acid metabolism” (adj p<0.05) (Fig. 1A **and Table S1B**). We identified other enriched pathways that contain both up-regulated and down-regulated genes, such as “phosphorylation” and “skeletal muscle cell differentiation”, as well as pathways that were enriched in down-regulated genes, such as “Response to interferon-beta”. (Fig. 1A **and Table S1B**). In summary, we unveiled a transcriptional signature in LCAD KO hearts characterized by the up-regulation of genes involved in amino acid metabolism, carboxylic acid metabolism, and ribosomal RNA metabolism.

**Figure 1.**
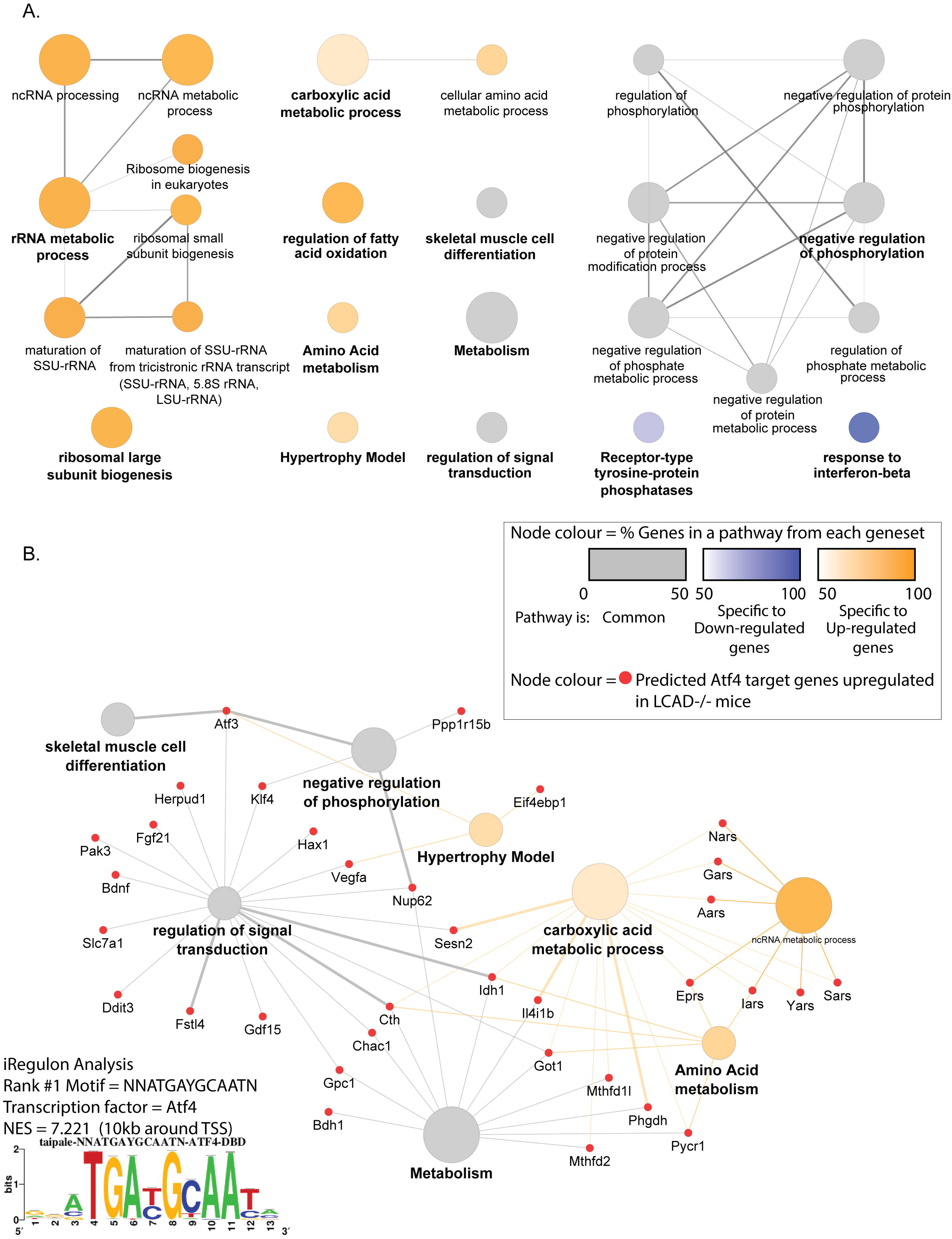
Pathway enrichment and transcription factor binding motif enrichment analysis of LCAD KO heart transcriptomic data. **A)** Pathway enrichment analysis of the DEGs between hearts from WT and LCAD KO mice according to GO (BP), KEGG, Wiki and Reactome databases. Genes that were either up- or down-regulated (logFC > 0.58, adjP < 0.05) were queried as two separated gene clusters. Network node annotations are the pathway terms found significantly enriched at adj P < 0.05, with size of node reflecting term significance, with the bigger nodes the more significant ones. The node color shows the proportion of genes from each cluster that are associated with the term. Grey nodes indicate genes associated to a term are from both the up- and down clusters and thus common to both gene lists. Nodes colored shades of orange or blue indicate genes that associate to a term (at least >60%) are more specific to either the up- or down-regulated geneset, respectively. Edges represent the ClueGO kappa score, which defines term-term interrelations based on shared genes between the terms. Terms are grouped at default kappa > 0.4) with edge thickness reflecting greater kappa value. Full results are in Table S1B. **B)** iRegulon transcription factor analysis was performed on the up-regulated differentially expressed genes using their putative regulatory regions 10kb centered around the transcription start site. The top ranked enriched motif had a significant normalized enrichment score (NES) of 7.221 and its sequence is shown in the figure. The transcription factor associated with the motif was ATF4. Fifty-two ATF4-target genes were found on the up-regulated geneset, a subset of those (red nodes) genes are shown according to the biological pathways they were found associated with in panel A. Full results are in Table S1C.

We next focused on transcriptional regulators that could explain the observed changes in gene expression and performed cis-regulatory sequence analysis. We found that the LCAD KO heart transcriptional signature was enriched for genes containing binding motifs that mapped for the transcription factor ATF4 (Fig. 1B **and Table S1C**). ATF4 is a key mediator of the ISR, a well conserved homeostatic response that reprograms translation in response to nutrient stress (Costa-Mattioli & Walter 2020; Harding et al. 2000; Lu et al. 2004; Quirós et al. 2017). Asparagine synthetase (*Asns*), the most upregulated gene in LCAD KO hearts, is a canonical ATF4 target gene (Siu et al. 2002). We found 51 additional ATF4 target genes in the up-regulated geneset (Fig. 1B **and Table S1C**). These results suggest that ATF4 is a key mediator of the transcriptional signature of hearts from LCAD KO mice after food withdrawal.

### Activation of the integrated stress response in the heart of LCAD KO mice

Next, we studied the ISR at the protein level using immunoblot in heart samples from LCAD KO mice isolated after overnight food withdrawal. When compared to controls, LCAD KO mice had increased cardiac protein levels of phosphorylated eIF2α (S51), ATF4 and ASNS (Fig 2A), which is consistent with activation of the ISR. We also found a decreased Thr37/46 phosphorylation of the eukaryotic translation initiation factor 4E-binding protein 1 (4E-BP1) in LCAD KO hearts compared with WT hearts (Fig 2A). 4E-BP1 inhibits protein translation by preventing eIF4E to bind cap structures on mRNAs (Shu et al. 2020). 4E-BP1 (*Eif4ebp1*) is also an ATF4 target gene (Torrence et al. 2021) and, consistently, its expression was up-regulated in LCAD KO hearts (**Table S1A**). We also found that the autophagy markers p62 and LC3B-II were augmented in LCAD KO hearts compared with WT hearts (**Fig S1A**). The cardiac increase of LC3B-II protein was validated in an independent set of WT and LCAD KO samples (**Fig. S1B**). Thus, LCAD deficiency leads to nutrient stress and, as a consequence, activation of the ISR via the eIF2α/ATF4 axis.

**Figure 2.**
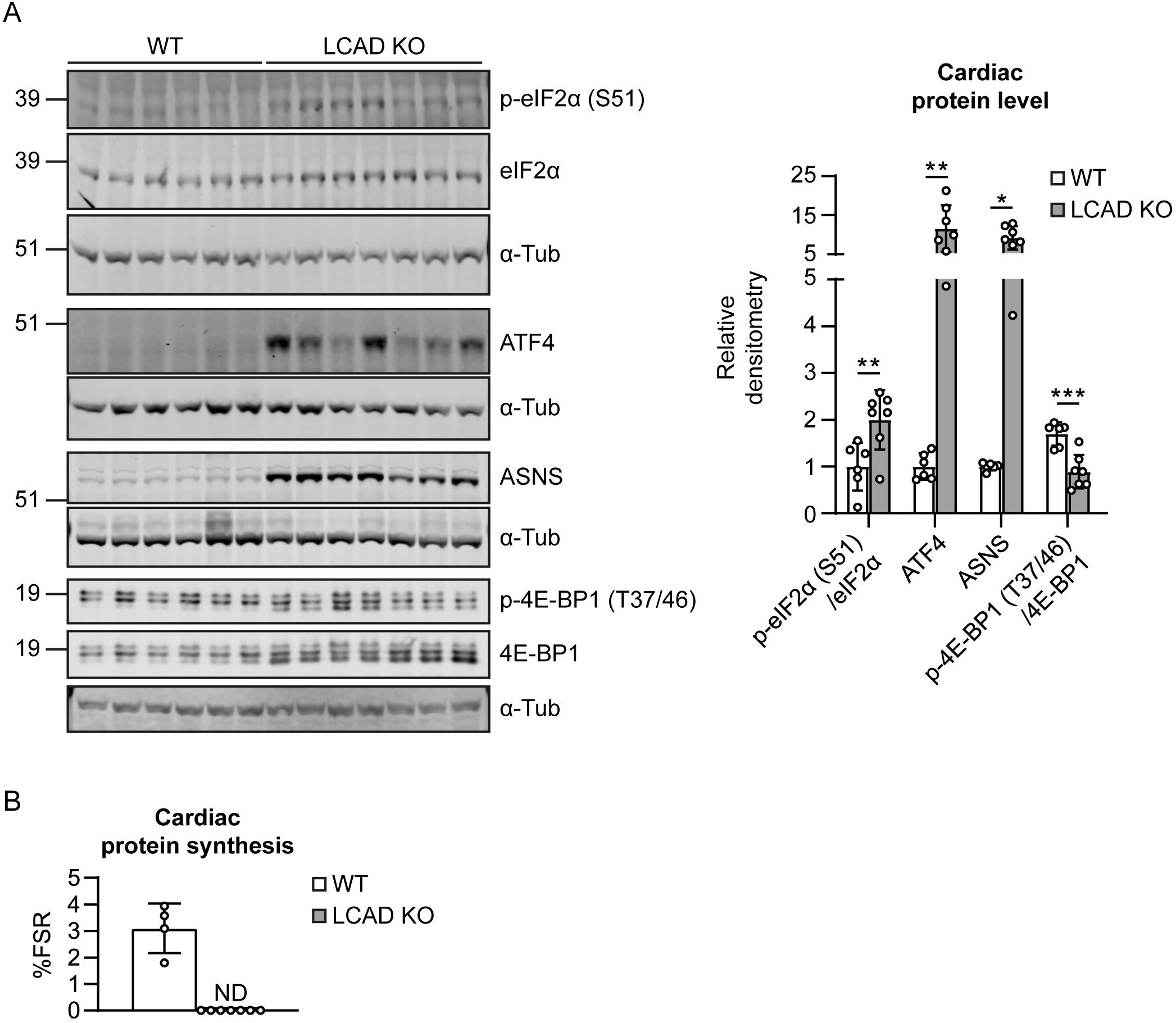
Activation of the integrated stress response in the heart of LCAD KO mice. **A**) Immunoblot of ISR markers p-eIF2α (S51), eIF2α, ATF4, ASNS, p-4E-BP1 (T37/46), 4E-BP1. Protein levels were quantified relative to α-tubulin (α-tub), or their non-phosphorylated form (as indicated on the x-axis label). **B**) Protein synthesis rate (FSR: Fractional synthesis rate, in %) was determined in the hearts of WT (n=4) and LCAD KO (n=7) mice after the injection of ^2^H_2_O and 8 hours of food withdrawal, by measuring the ^2^H-labeling of protein-bound alanine. ND: not detected. Individual values, the mean and the standard deviation are graphed. Statistical significance was tested using unpaired t test with Welch’s correction. * p<0.05; ** p<0.01; *** p<0.001

We measured protein synthesis rates in WT and LCAD KO hearts by the ^2^H_2_O method. WT hearts showed detectable protein synthesis during 8 hours of food withdrawal starting at the onset of the active period (dark cycle). In contrast, protein synthesis was not detectable in any of the in LCAD KO heart samples (Fig. 2B). No differences in protein synthesis were detected during the light cycle (**Fig. S1C**). Thus decreased nutrient availability can lead to a pronounced reduction in protein synthesis in the heart, which is likely mediated by the ISR.

### Metabolic alterations in response to food availability in LCAD KO mice

Overnight food withdrawal leads to significant changes in mouse metabolism and pronounced differences between WT and LCAD KO mice. In order to identify early and late changes in response to food withdrawal in the LCAD KO mouse model, we collected plasma and heart samples from WT and LCAD KO mice every 4 hours during a 12-hour food withdrawal period. Food was withdrawn just before the dark phase at 6 p.m. and a random fed sample was collected (time point 1). During food withdrawal, samples were collected at 10 p.m., 2 a.m. and 6 a.m. (time points 2, 3 and 4). At 6 a.m. mice were refed for 4-hours with sample collection at 10 a.m. (time point 5). Additionally, we collected samples from animals that were randomly fed and sacrificed at 10 a.m. (time point 6). Analysis of plasma metabolites (**Table S2A, S2B**) showed that many of the differences in circulating metabolites between the LCAD KO and WT mice were already evident 4 hours after food withdrawal including a peak in C14:1-carnitine accumulation (Fig. 3A), free carnitine deficiency (Fig. 3B), hypoketosis (Fig. 3C, assessed by C4OH-carnitine levels) (Soeters et al. 2012), and hypoglycemia (Fig. 3D). Some of the differences between LCAD KO and WT animals became more pronounced with time (carnitine deficiency), whereas others stabilized (hypoglycemia) (Fig. 3B, D). A period of 4 hours refeeding normalized blood glucose (Fig. 3D), but free carnitine concentration remained low (Fig. 3B). Accumulation of long-chain acylcarnitines in plasma was evident at every time point during fasting, but decreased sharply upon refeeding (Fig. 3A, **S2A, and Table S2A**).

**Figure 3.**
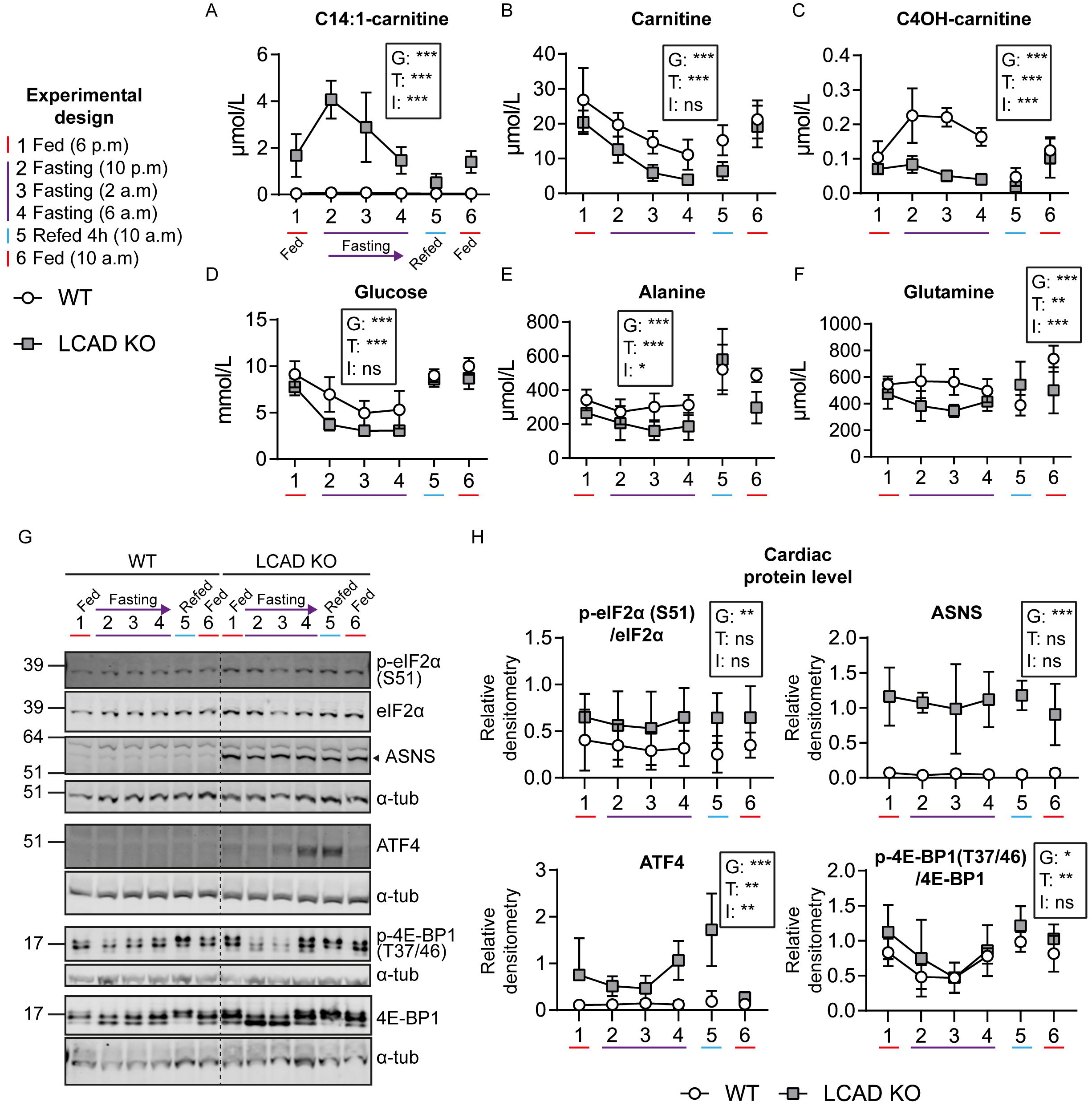
Metabolic alterations in response to food availability in plasma and heart of WT and LCAD KO mice. **(A-F)** Effect of food availability on plasma metabolite levels at different time points in WT (n=6 per time point) and LCAD KO (n=5-6 per time point) mice. Time point 1: 6 p.m. (random fed), 2: 10 p.m. (4 hours after food withdrawal), 3: 2 a.m. (8 hours after food withdrawal), 4: 6 a.m. (12 hours after food withdrawal), 5: 6 a.m. (4 hours of refeeding after 12 hours of food withdrawal), 6: 10 a.m. (random fed). **A**) C14:1-carnitine, **B**) carnitine, **C**) C4OH-carnitine, **D**) glucose, **E**) alanine, **F**) glutamine. **G**) Representative immunoblots of ISR markers in WT and LCAD KO hearts at different time points of the food withdrawal time course study (n=4 per genotype and time point) **H**) Band densitometry was quantified relative to α-tub, or the non-phosphorylated form of the protein. Individual values, the mean and the standard deviation are represented. The result of the two-way ANOVA test is displayed within the graphs. G: “Genotype”. T: “Time point”. I: “Interaction”. * p<0.05; ** p<0.01; *** p<0.001

The reduction in the plasma level of gluconeogenic amino acids such as alanine and glutamine was most pronounced after 8 hours of fasting but also present 4 hours after food withdrawal (Fig. 3E, F). Notably, levels of alanine and glutamine were only comparable to WT upon refeeding (time point 5), suggesting that the reduced availability of several amino acids is already noticeable after a short period of without feeding (Fig. 3E, F). A similar profile was found for other amino acids such as glutamate, aspartate and proline (**Fig. S2A and Table S2B**). Plasma levels of serine and glycine increased considerably after refeeding in LCAD KO mice (**Fig. S2A**). In contrast, the contribution of branched-chain amino acids (BCAA) to the total pool of amino acids increased 4 hours after food withdrawal, remained elevated during the period of food withdrawal, and returned to basal levels after refeeding (**Fig. S2A**). Together, these data show that the metabolic phenotype of LCAD KO mice characterized by hypoketotic hypoglycemia, increased levels of C14:1-carnitine, secondary carnitine deficiency, and reduced amino acid availability, is present early after food withdrawal.

The same samples were used to study early and late events in metabolic signaling in the heart of LCAD KO mice. We found that the levels of phospho-eIF2α, ATF4 and ASNS, indicative of activation of the ISR, were increased at all tested time points in LCAD KO hearts compared with WT hearts (Fig. 3G, H). ATF4 protein levels reached their peak after refeeding (Fig. 3G, H). Phosphorylated 4E-BP1 (T37/46) progressively decreased with time in WT and LCAD KO hearts and drastically increased after refeeding in both genotypes (Fig. 3G, H), in line with mTORC1 activation after refeeding (Ben-Sahra et al. 2016; Torrence et al. 2021). LC3B-II was increased in LCAD KO hearts at time points 1 to 4, then decreased upon refeeding (**3G, H**), consistent with an inhibition of autophagy after feeding. Levels of p62 were increased in the heart of refed LCAD KO mice. However, no difference was observed after food was withdrawn (time points 2, 3 and 4) (**Fig. S2B**). These results indicate that the activity of the ISR is higher in LCAD KO hearts when compared to WT hearts irrespective of the feeding status of the mice.

### Accumulation of uncharged tRNAs in the LCAD KO mouse heart

Four different kinases (GCN2, PERK, HRI, and PKR) sense different types of nutrient stress and converge on eIF2α phosphorylation to initiate the ISR (Costa-Mattioli & Walter 2020). We did not find any signs of PERK (encoded by *Eif2ak3*) activation, an eIF2α kinase that senses protein folding alterations in the endoplasmic reticulum (Harding et al. 1999, 2003). Protein levels of ATF6, phosphorylated PERK (Thr980), or the ER chaperone BiP, which is also referred to as GRP78 or HSPA5, did not change in LCAD KO hearts compared with WT hearts (**Fig. S3A**). Moreover, we did not find any change in XBP1 splicing events (Yoshida et al. 2001) in LCAD KO hearts compared with WT hearts (**Fig. S3B**). Hence, we found no evidence that ER stress acts as the activator of the ISR in LCAD KO hearts.

We hypothesized that the most likely mediator of eIF2α phosphorylation in LCAD KO hearts is the general control nonderepressible 2 (GCN2) kinase because this kinase is activated by reduced amino acid availability (Berlanga et al. 1999; Inglis et al. 2019; Ishimura et al. 2016; Pavlova et al. 2020; Wek et al. 1995; Zhang et al. 2002). GCN2 is autophosphorylated in the presence of uncharged transfer RNA (tRNA) (Dong et al. 2000; Wek et al. 1995), which may accumulate due to decreased availability of free amino acids (Pavlova et al. 2020). We have previously described how an impairment in long-chain fatty acid oxidation leads to a depletion in the levels of free amino acids in plasma and tissues (Houten et al. 2013; Ranea-Robles et al. 2020) and a defect in cardiac anaplerosis (Bakermans et al. 2013). With the available antibodies, however, we were unable to detect phosphorylated GCN2 in mouse heart samples (not shown).

To assess the potential activators of GCN2, we measured tRNA charging in heart samples from WT and LCAD KO hearts after a short period of 4 hours of food withdrawal (Pavlova et al. 2020). We found significant decreases in the charging of tRNA^Glu^, tRNA^Gln^, tRNA^Pro^, tRNA^Arg^ tRNA^Leu^, and tRNA^Val^ (Fig. 4A **and S4A**). Amino acid charging of tRNA^eMet^ tended to decrease in LCAD KO hearts (Fig. 4A). tRNA^Asp^ charging tended to increase, whereas tRNA^Ala^ tended to decrease, only in male LCAD KO hearts (**Fig. S4A**). We measured amino acid concentrations in the same heart samples and found decreased levels of several amino acids including glutamate, glutamine, methionine, histidine and citrulline in LCAD KO hearts when compared to WT hearts (Fig. 4B **and S4B, and Table S3**). The decreased levels of Glu, Gln, and Met parallel the decrease in charging of their cognate tRNA (Fig. 4A and 4B). The concentration of several other amino acids such as proline, glycine, and lysine increased in LCAD KO hearts (Fig. 4B **and S4B, and Table S3**). Remarkably, tRNA^Pro^ was decreased in these samples (Fig. 4A). These results show that, after a short period of food withdrawal, a defect in long-chain FAO leads to pronounced changes in amino acid availability and the selective uncharging of several tRNAs in the heart. This finding suggests that GCN2 kinase is the likely effector of the activation of the ISR in LCAD KO hearts.

**Figure 4.**
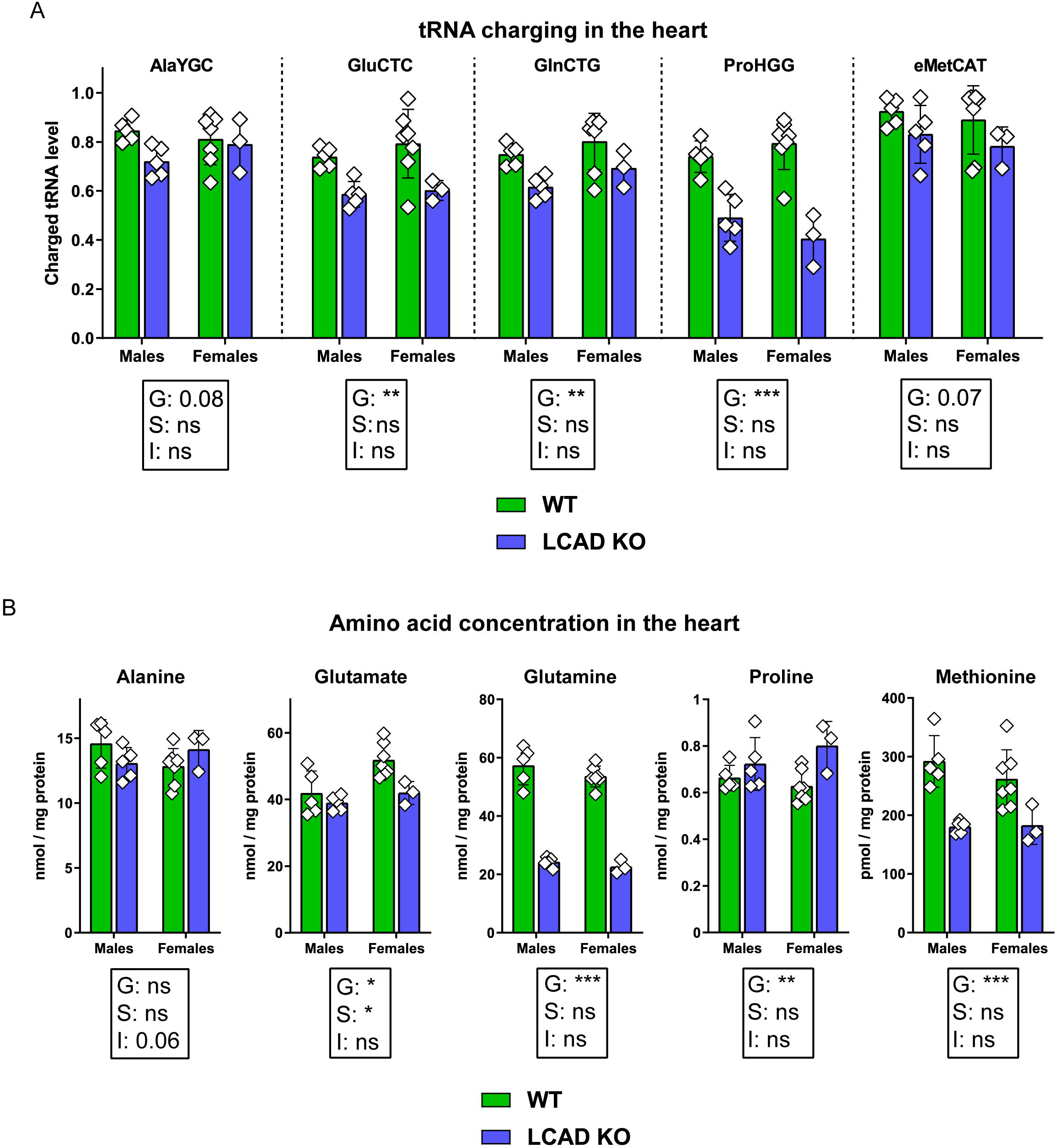
tRNA charging and amino acid concentrations in mouse heart after a short period of food withdrawal. **A**) tRNA charging of the following isodecoder groups was measured in mouse heart using specific primers: AlaYGC, GluCTC, GlnCTG, ProHGG, and eMetCAT. The last three letters denote an anticodon or a group of anticodons targeted. Y: C or T; H: A, C or T. **B**) Amino acid concentrations (in pmol/mg protein or nmol/mg protein) were measured in mouse heart. N = 3-7 biological replicates per genotype (WT and LCAD KO) and sex (males and females). Individual values, the mean and the standard deviation are represented. The result of the two-way ANOVA test is displayed within the graphs. G: “Genotype”. S: “Sex”. I: “Interaction”. * p<0.05; ** p<0.01; *** p<0.001

### Activation of the ISR in CPT2-deficient mouse hearts

We decided to investigate if the activation of the ISR could be observed in other models of long-chain FAO defects. We first compared the transcriptional signature of LCAD KO hearts to the signature from hearts of cardiac and skeletal muscle-specific CPT2 KO (*Cpt2*^M-/-^) mice (Pereyra et al. 2017). There was a significant overlap between the up- and down-regulated DEGs from these two models with a defect in long-chain FAO (Fig. 5A, 5B, and **Table S4A**). Importantly, the shared upregulated genes were enriched for key terms such as “Amino-tRNA biosynthesis”, “Amino acid metabolism”, and “Amino acid transport across the plasma membrane” **(Fig. S5 and Table S4B)**. This shared transcriptional signature from the hearts of two models of long-chain FAO defects was enriched for genes containing binding motifs that mapped for the transcription factors ATF4 **(Fig. S5 and Table S4C)**. These results suggest that activation of the ISR/ATF4 axis and disturbances in amino acid metabolism are common adaptations of the long-chain FAO deficient heart.

**Figure 5.**
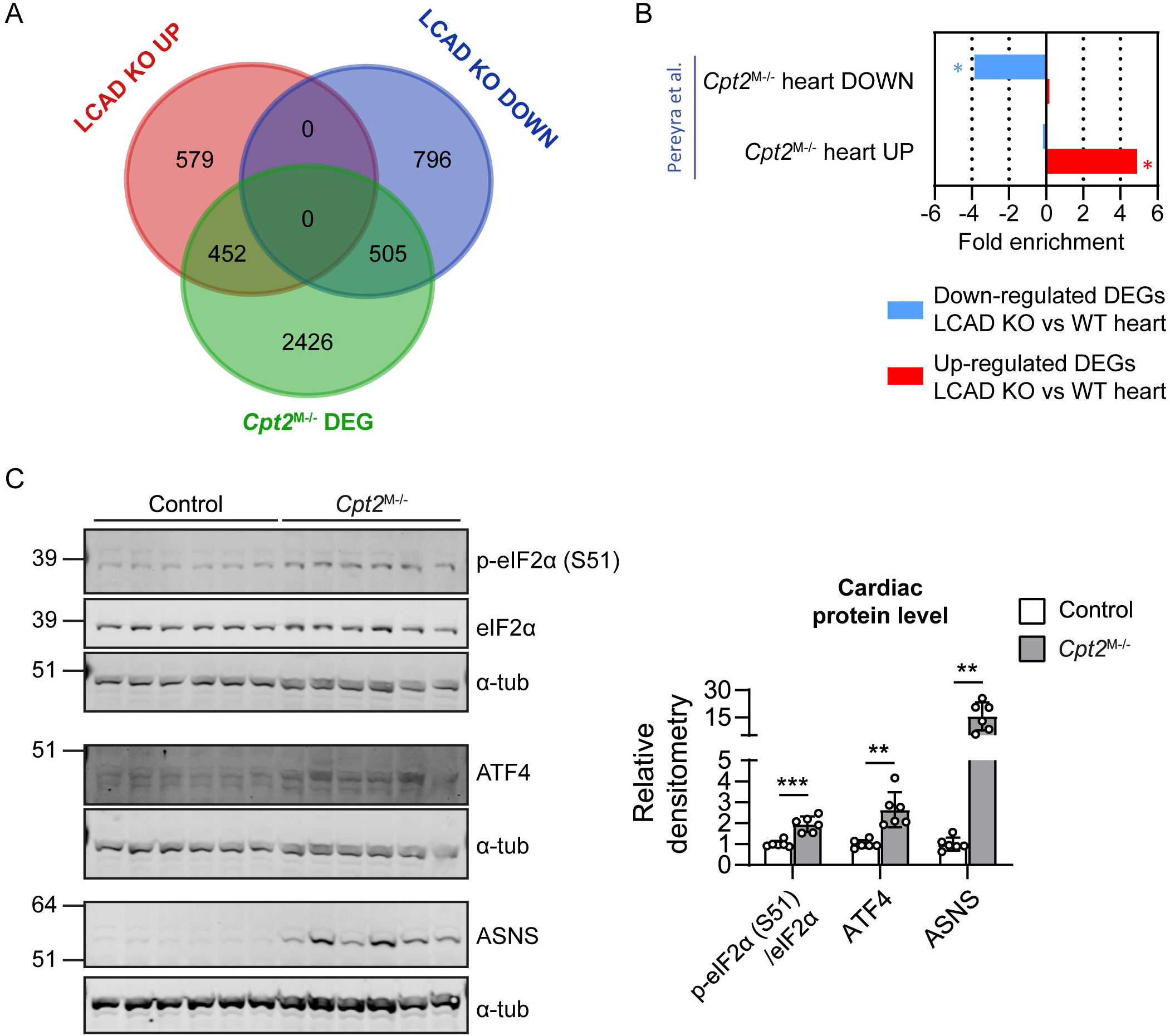
Cardiac ISR in other long-chain FAO deficient models. **A**) Venn diagrams comparing the number of DEGs obtained from LCAD KO hearts with the number of DEGs obtained from *Cpt2*^M-/-^ hearts (Pereyra et al. 2017). **B**) Results of enrichment analysis of genes either up- or down-regulated in LCAD KO hearts in two gene sets curated from the transcriptional signature of *Cpt2*^M-/-^ hearts (Pereyra et al. 2017). Values represent the fold enrichment and significance is indicated as * (adj p<0.001). Full table of results are in Table S4A. **C**) Immunoblots of the ISR markers phosphorylated eIF2α (p-eIF2α) at S51, ATF4 and ASNS in the hearts of control and CPT2-deficient (*Cpt2*^M-/-^) mice (n=6) are shown. Band densitometry was quantified relative to α-tub, or the non-phosphorylated form of the protein, as indicated. Individual values, the average and the standard deviation are graphed. The result of the unpaired t test with Welch’s correction is displayed within the graphs. ** p<0.01; *** p<0.001.

We then explored whether the activation of the ISR was also present at the protein level of the CPT2-deficient hearts. We observed an activation of the ISR in *Cpt2*^M-/-^ hearts, reflected by increased levels of phosphorylated eIF2α (S51), ATF4, and ASNS (Fig. 5C), similar to what we observed in the LCAD KO hearts. Together, these results indicate that the cardiac activation of the ISR is a conserved mechanism in long-chain FAO defects.

## Discussion

In this study, we assessed the effects of nutrient availability and a long-chain FAO defect on mRNA expression and metabolic signaling in the heart. We show that the nutrient stress caused by LCAD deficiency activates the ISR and impacts cardiac protein synthesis and amino acid metabolism. We show here for the first time that a defect in long-chain FAO can lead to the selective uncharging of several tRNAs, in line with the reduction in the intracellular free amino acid levels reported in LCAD KO hearts (Houten et al. 2013; Ranea-Robles et al. 2020). The upregulation of the tRNA aminoacylation pathway in the LCAD KO and the CPT2 KO hearts supports these findings and suggests it may represent a common pathophysiological mechanism in long-chain FAO defects. This is not only the first report on tRNA charging in the mouse heart, this is also the first time that uncharging of tRNAs is demonstrated in a disease model.

The activation of the ISR may serve to attenuate global mRNA translation and conserve amino acids and energy, as one adaptive response to overcome the nutritional stress imposed by the long-chain FAO defect. The activation of ISR was independent of feeding status, which may be explained by the fact that the heart typically relies on FAO for the majority of its ATP production. These findings may be particularly relevant for patients with long-chain FAO defects that are typically under dietary management to prevent a metabolic crisis due to increased reliance on FAO. Moreover, these findings suggest that ISR activation and changes in amino acid metabolism and tRNA charging may be involved in the pathophysiology of pathologic cardiac hypertrophy, which is characterized by a decreased fatty acid metabolism.

eIF2α can be phosphorylated by PERK, GCN2, PKR and HRI kinases, which are activated by unfolded proteins in the ER (Harding et al. 1999, 2000), amino acid deficiency (Berlanga et al. 1999; Harding et al. 2000, 2003; Ishimura et al. 2016; Wek et al. 1995; Zhang et al. 2002), viral infection and heme deficiency (Costa-Mattioli & Walter 2020), respectively. Activation of the ISR after a short period of food withdrawal (only 4 hours) was concomitant with changes in the availability of many amino acids and the selective uncharging of certain tRNAs in LCAD KO hearts. Activation of the ISR in LCAD KO hearts was also associated with a reduction in protein synthesis. Previously, decreased amino acid availability was demonstrated to reduce the charging of tRNA^Gln^ *in vitro* (Pavlova et al. 2020). Thus, amino acid limitation-associated uncharging of tRNAs, and in particular tRNA^Gln^, may cause the induction of the ISR in LCAD KO hearts via GCN2 kinase activation. Other studies have confirmed the role of GCN2 kinase in modulating homeostatic responses in the heart (Feng et al. 2019; Lu et al. 2014). We cannot completely exclude a contribution of other eIF2α kinases to the ISR activation in this model. These include HRI and PKR, which have been recently linked to the sensing of mitochondrial-derived stress (Condon et al. 2021; Fessler et al. 2020; Guo et al. 2020; Kim et al. 2018). Genetic experiments with mouse models deficient for each of these kinases or pharmacological modulation of eIF2α phosphorylation should help to determine the mechanism of eIF2α activation in long-chain FAO deficiency and whether this ISR activation is a protective or a maladaptive response of the FAO-deficient heart.

We show here that ATF4 transcription factor binding motif is enriched in the DEGs of LCAD KO hearts. ISR and ATF4 activation is a common finding in mitochondrial dysfunction caused by different mitochondrial stressors, such as cardiac-specific mitochondrial tRNA synthetase deficiency (Dogan et al. 2014), mtDNA gene expression defects (Forsström et al. 2019; Kühl et al. 2017), electron transport chain inhibition (Mick et al. 2020), and mitochondrial translation inhibition, mitochondrial membrane potential disruption, and mitochondrial protein import inhibition (Quirós et al. 2017), but has not been studied in models for FAO deficiency. Our pathway enrichment analysis using the shared up-regulated genes from the heart of LCAD KO and muscle-specific CPT2-deficient mice (Pereyra et al. 2017) unveiled an activation of amino acid metabolism pathways and an induction of the transcription factor ATF4, which we subsequently validated at the protein level. This suggests that long-chain FAO defects impact on cardiac amino acid metabolism and activate the ISR.

The activation of the ISR in the heart of LCAD KO mice appears largely independent of the feeding status, which may suggest that it plays a role in the development of the observed cardiac hypertrophy. LCAD KO mice display a non-progressive cardiac hypertrophy (Bakermans et al. 2011), but cardiac dysfunction only arises upon food withdrawal (Bakermans et al. 2011, 2013). Indeed, some of the characteristic features of the ISR are associated with an adaptive responsive that may explain cardiac hypertrophy in LCAD KO mice. For example, during the ISR there is a redirection of glucose metabolic flux towards serine biosynthesis, which provides precursors for one-carbon (1C) metabolism, phospholipids and glutathione synthesis, as a protective mechanism for managing mitochondrial bioenergetic stress (Bao et al. 2016; Forsström et al. 2019; Gibb & Hill 2018; Nikkanen et al. 2016). In a recent metabolic profiling that we performed in overnight fasted LCAD KO hearts we found increased levels of asparagine, serine, glycine, pyrimidine nucleotides, and phospholipids in LCAD KO hearts (Ranea-Robles et al. 2020) consistent with the ATF4-mediated induction of the serine biosynthetic genes *Psat1* (Phosphoserine aminotransferase) and *Phgdh* (Phosphoglycerate dehydrogenase), and *Asns*. Here, we studied amino acid metabolism after a shorter period of food withdrawal, which is likely more translatable to human FAO disorders, and describe similarly profound changes in amino acid metabolism in LCAD KO hearts, including increased levels of proline, glycine, and lysine, and decreased levels of glutamine and glutamate. Based on the recent discovery of a biosynthetic arm downstream of ATF4 activation (Torrence et al. 2021), we speculate that the activation of the ISR and ATF4 in LCAD KO hearts may contribute to cardiac hypertrophy. Further investigations manipulating these metabolic adaptations in LCAD KO hearts are required to understand their contribution to cardiac hypertrophy and, ultimately, to nutrient stress-induced cardiac dysfunction in FAO defects.

In conclusion, we describe the progression of metabolic alterations in the plasma and the heart of LCAD KO mice during an overnight fasting, confirming that most of the metabolic changes start early after food withdrawal. We also report the selective uncharging of certain tRNAs in the heart of LCAD KO mice after 4 hours of food withdrawal. Collectively, these findings exemplify how defects in FAO remodels cardiac metabolism impacting on glucose and amino acid metabolism. The results obtained here provide a basis for future investigations to assess the impact of therapeutic modulations of the ISR on cardiac function in FAO deficiencies.

## Acknowledgements

The authors thank Dr. Tetyana Dodatko for her excellent technical assistance. We acknowledge the help of the shared resource facilities at the Icahn School of Medicine at Mount Sinai (Colony Management, Genomics Core and the Comparative Pathology Laboratory).

Research reported in this publication was supported by the National Institute of Diabetes and Digestive and Kidney Diseases of the National Institutes of Health under Award Number R01DK113172. The content is solely the responsibility of the authors and does not necessarily represent the official views of the National Institutes of Health.

