## Supplementary Material for "A mitochondrial long-chain fatty acid oxidation defect leads to uncharged tRNA accumulation and activation of the integrated stress response in the mouse heart"

### Supplemental information Table of Contents

**Supplementary Figure legend**

**Supplementary Figure S1.** Activation of the integrated stress response in the heart of LCAD KO mice

**Supplementary Figure S2.** Metabolic alterations in response to food availability in plasma and heart of WT and LCAD KO mice

**Supplementary Figure S3.** Activation of the integrated stress response in the heart of LCAD KO mice

**Supplementary Figure S4.** tRNA charging and amino acid concentrations in mouse heart after a short period of food withdrawal

**Supplementary Figure S5.** Pathway enrichment and transcription factor binding motif enrichment analysis of the shared up-regulated genes between the LCAD KO heart and the *Cpt2*$^{M-/-}$ heart transcriptomic data

**Supplementary Tables**

**Supplementary Table S1.** LCAD KO heart RNA-seq signature

**Supplementary Table S2.** Plasma metabolic profile of WT and LCAD KO mice during the food withdrawal time-course experiment

**Supplementary Table S3.** Cardiac amino acid profile of WT and LCAD KO mice fasted during 4 hours.

**Supplementary Table S4.** Cardiac amino acid profile of WT and LCAD KO mice fasted during 4 hours.

**Supplementary Table S5.** Comparison of LCAD KO vs CPT2 KO heart DEGs.

### Supplementary Figure legend

**Supplementary Figure S1**. Activation of the integrated stress response in the heart of LCAD KO mice.

**A**) Immunoblot of autophagy markers p62 and LC3B. Protein levels were quantified relative to alpha-tubulin (α-Tub). **B**) Immunoblot of autophagy marker LC3B in an independent set of WT and LCAD KO hearts (n=5). α-tub was used as loading control. **C**) Protein synthesis rate (FSR: Fractional synthesis rate, in %) was determined in the hearts of two sets of WT (n=4 per set) and LCAD KO (n=4 per set) mice after the injection of $^{2}H_2O$. Mice were then either fed *ad libitum* for 7 hours (left panel) or food was removed during 7 hours during the morning (right panel). FSR was determined by measuring the $^{2}$H-labeling of protein-bound alanine. Individual values, the mean and the standard deviation are graphed (A, C). Statistical significance was tested using unpaired t test with Welch's correction (A, C). * p<0.05; ** p<0.01; *** p<0.001

**Supplementary Figure S2**. Metabolic alterations in response to food availability in plasma and heart of WT and LCAD KO mice.

**A**) Effect of food availability on plasma metabolite levels (µmol/L or ratio in the case of BCAA/Total AA) at different time points in WT (n=6 per time point) and LCAD KO (n=5-6 per time point) mice. Time point 1: 6 p.m. (random fed), 2: 10 p.m. (4 hours after food withdrawal), 3: 2 a.m. (8 hours after food withdrawal), 4: 6 a.m. (12 hours after food withdrawal), 5: 10 a.m. (4 hours of refeeding after 12 hours of food withdrawal), 6: 10 a.m. (random fed). **B**) Effect of food availability on autophagy markers p62 and LC3B-II. Representative immunoblots are shown. Protein levels were quantified relative to α-Tub. The average and the standard deviation for each metabolite and protein are graphed. The result of the two-way ANOVA test is displayed within the graphs. G: Genotype. T: Time point. I: Interaction. * p<0.05; ** p<0.01; *** p<0.001

**Supplementary Figure S3**. Activation of the integrated stress response in the heart of LCAD KO mice

**A**) Immunoblot of ER stress markers. Last lane in pPERK (T980), PERK and ATF6 immunoblots has a positive control sample obtained from HEK293 cells treated 6 hours with 2 µg/mL tunicamycin. Protein levels were quantified relative to α-tub. **B**) Percentage of spliced *Xbp1* mRNA reads relative to the total *Xbp1* mRNA reads as measured by RNA-sequencing in WT and LCAD KO hearts. Individual values, the mean and the standard deviation are graphed. Statistical significance was tested using unpaired t test with Welch's correction.

**Supplementary Fig. S4**. tRNA charging and amino acid concentrations in mouse heart after a short period of food withdrawal

**A**) tRNA charging of the following isodecoder groups was measured in mouse heart using specific primers: AspGTC, ArgAGC, LeuWAG, and ValMAC. The last three letters denote an anticodon or a group of anticodons targeted. W: A or T; M: A or C. **B**) Amino acid concentrations (in pmol/mg protein or nmol/mg protein) were measured in mouse heart. N = 3-7 biological replicates per genotype (WT and LCAD KO) and sex (males and females). Individual values, the mean and the standard deviation are represented. The result of the two-way ANOVA test is displayed within the graphs. G: “Genotype”. S: “Sex”. I: “Interaction”. * $p<0.05$; ** $p<0.01$; *** $p<0.001$

**Supplementary Figure S5**. Pathway enrichment and transcription factor binding motif enrichment analysis of the shared up-regulated genes between the LCAD KO heart and the *Cpt2*$^{M-/-}$ heart transcriptomic data

**A)** Pathway enrichment analysis of the shared up-regulated DEGs between hearts from WT and LCAD KO mice, and hearts from WT and *Cpt2*$^{M-/-}$ mice (Pereyra et al. 2017), according to KEGG, Wiki and Reactome databases. Genes that were up-regulated (adjP < 0.05) from the full output of Pereyra et al. were queried to compare them with the LCAD KO heart transcriptional signature. Full table of results are in Table S4B. Network node annotations are the pathway terms found significantly enriched at adj P < 0.05, with size of node reflecting term significance, with the bigger nodes the more significant ones. Edges represent the ClueGO kappa score, which defines term-term interrelations based on shared genes between the terms. Terms are grouped at default kappa > 0.4, with edge thickness reflecting greater kappa value. iRegulon transcription factor analysis was performed on the shared up-regulated DEGs using their putative regulatory regions 10kb centered around the transcription start site. The top ranked enriched motif had a significant normalized enrichment score (NES) of 6.764 and its sequence is shown in the figure. The transcription factor associated with the motif was ATF4. Full results are in Table S4C.

**Supplementary Figure S1**

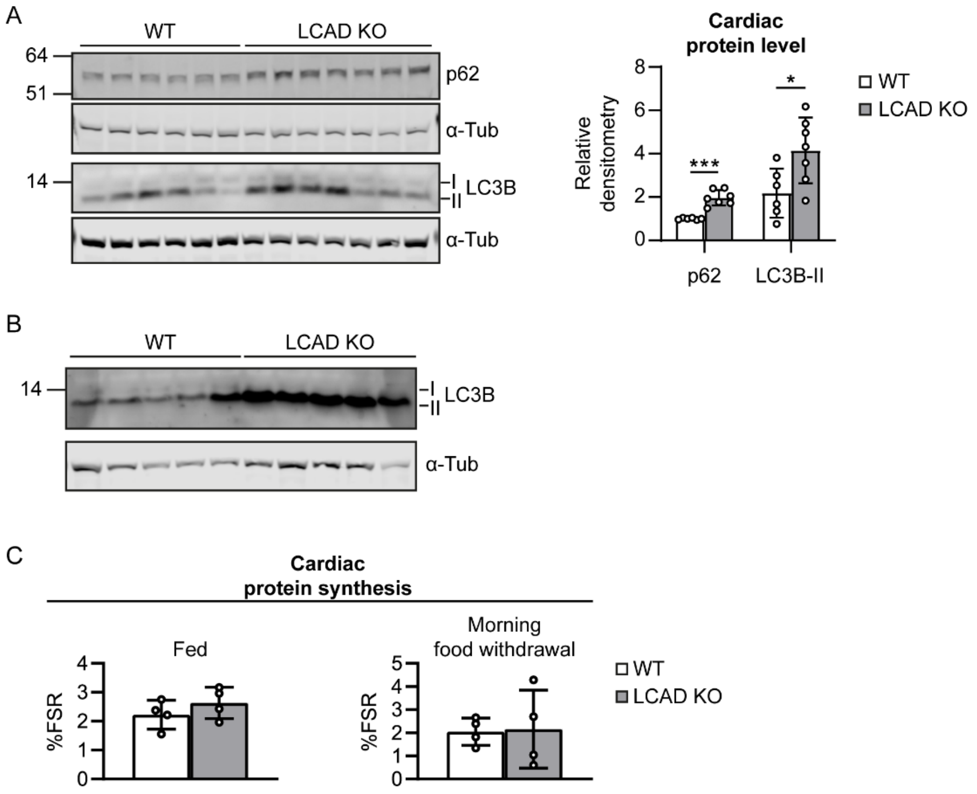

**Supplementary Figure S2**

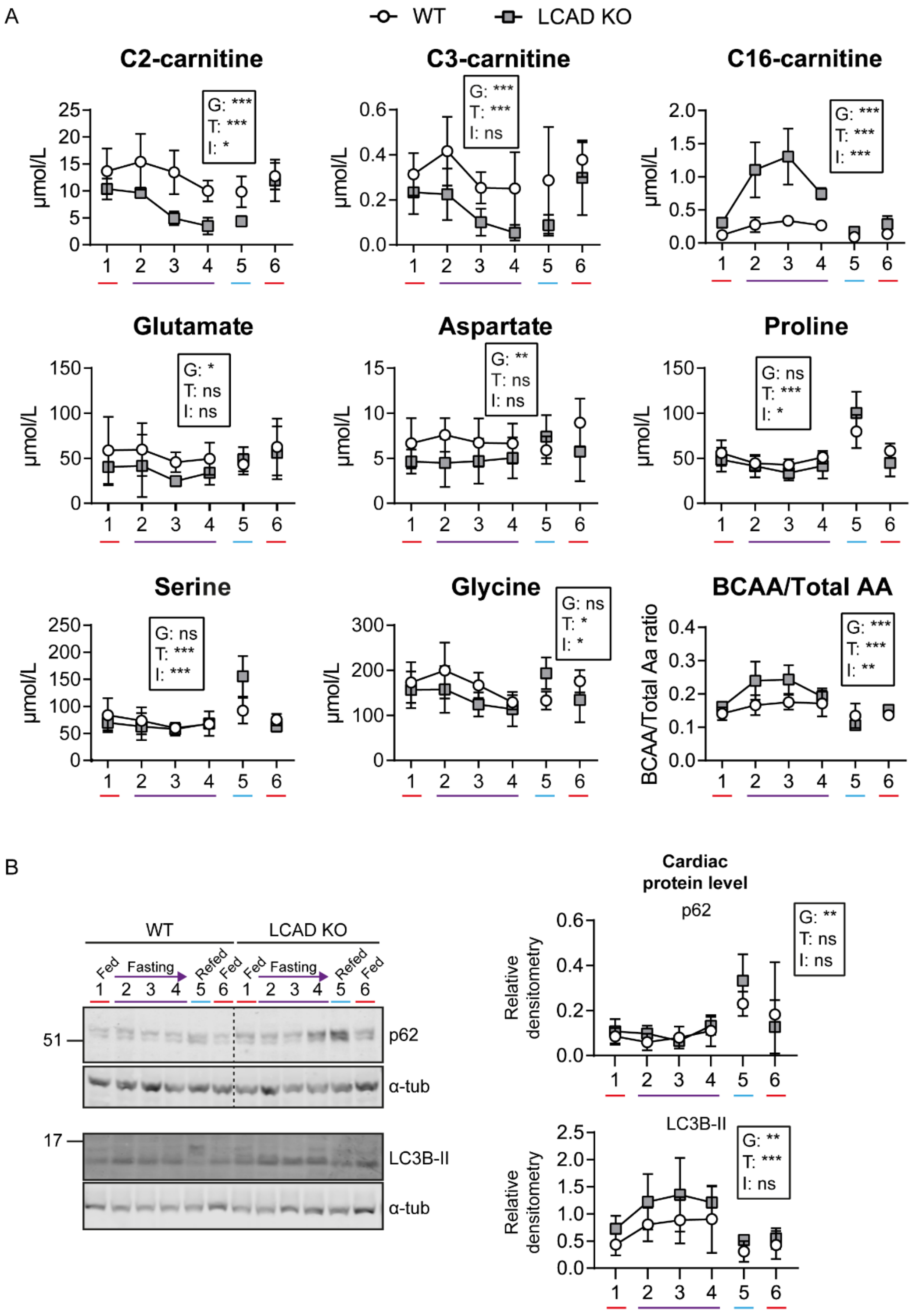

**Supplementary Figure S3**

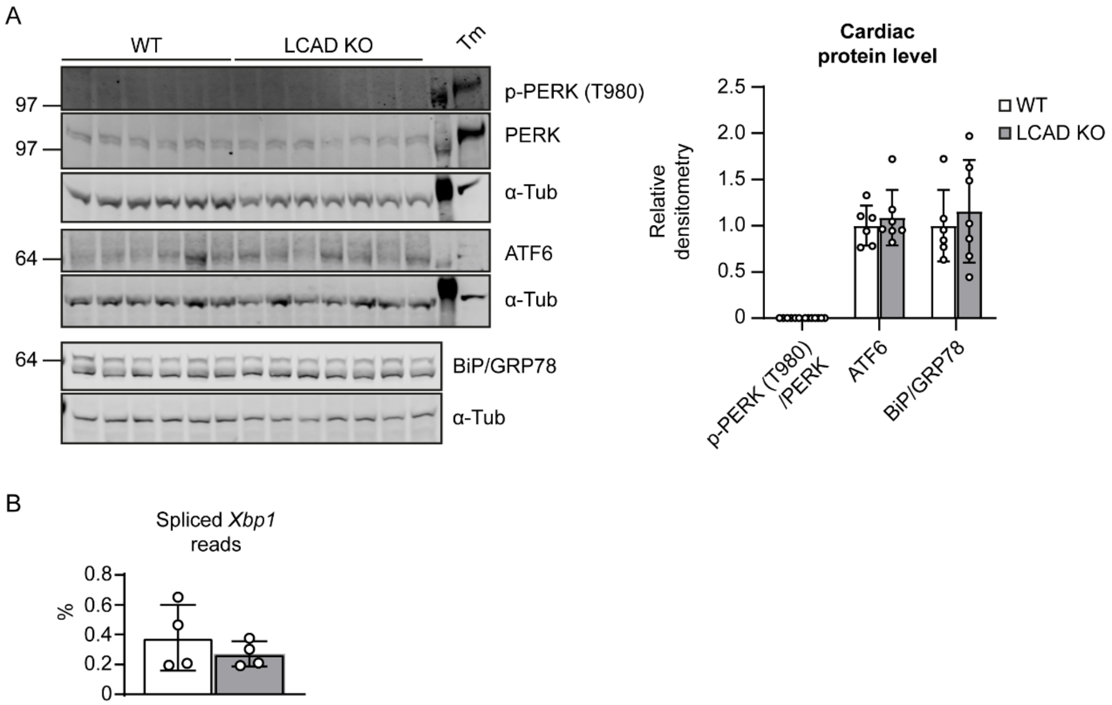

**Supplementary Figure S4**

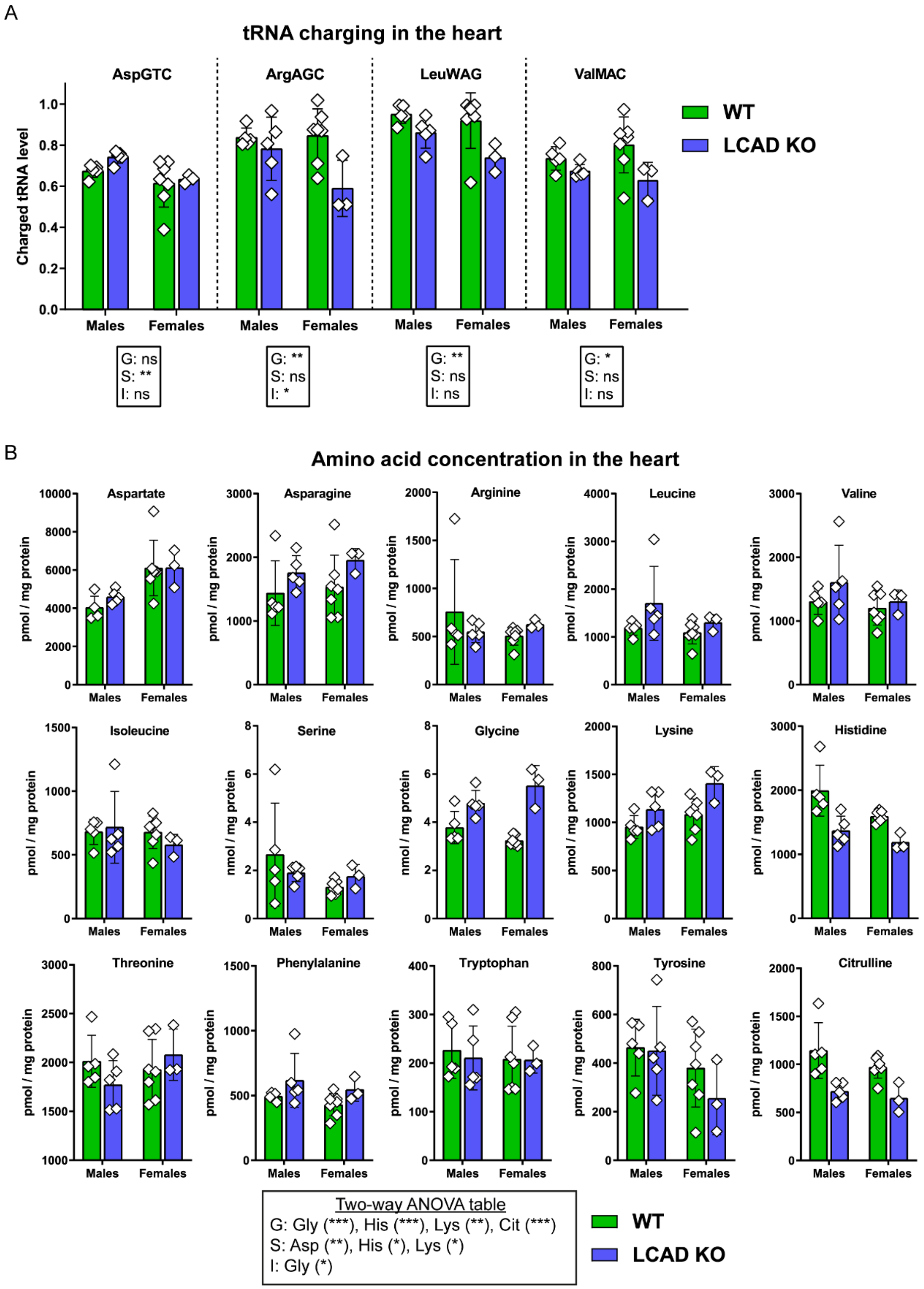

**Supplementary Figure S5**

A

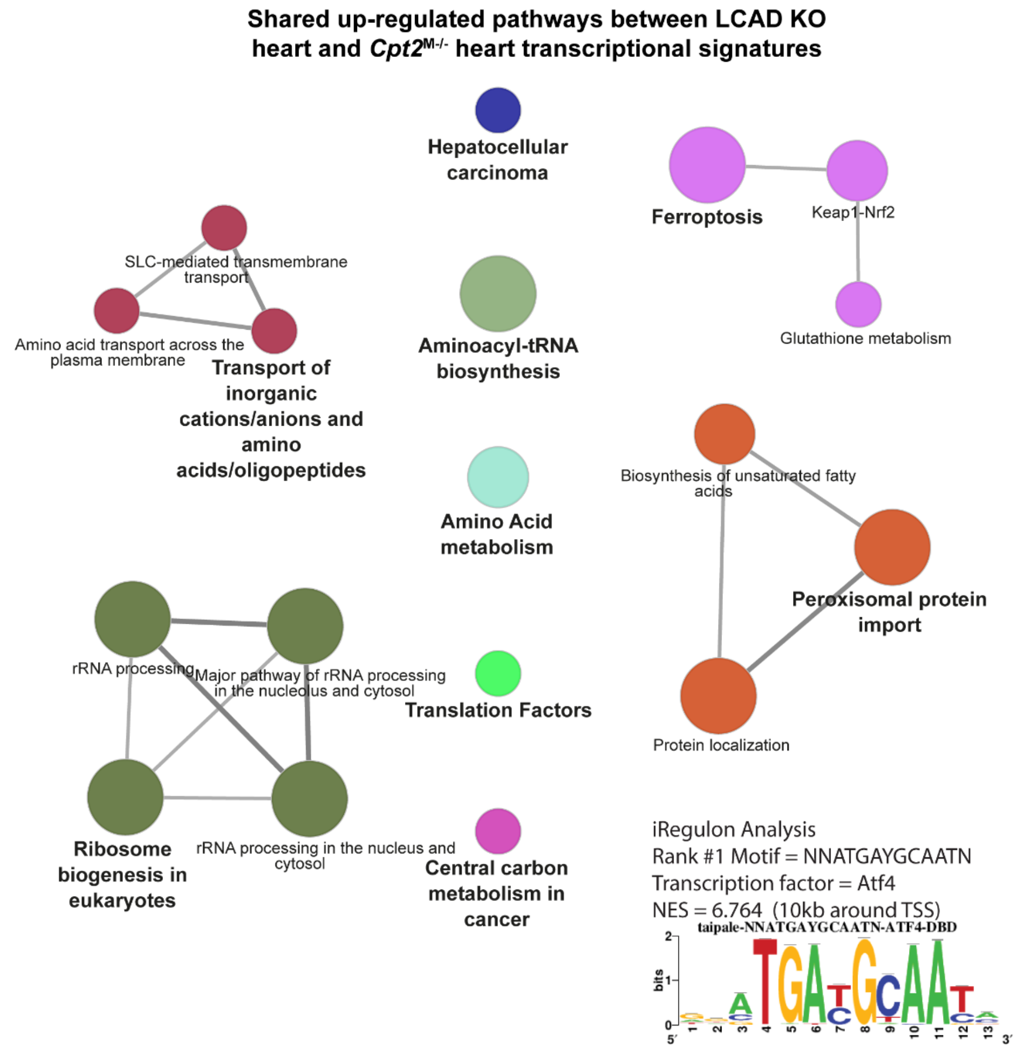
